# Machine learning versus logistic regression methods for 2-year mortality prognostication in a small, heterogeneous glioma database

**DOI:** 10.1101/472555

**Authors:** Sandip S Panesar, Rhett N D’Souza, Fang-Cheng Yeh, Juan C Fernandez-Miranda

**Author notes:** **Corresponding Author**: Sandip S Panesar MD MSc Department of Neurosurgery Stanford University 300 Pasteur Drive Palo Alto California 94025 United States of America.

## Abstract

**Background:** Machine learning (ML) is the application of specialized algorithms to datasets for trend delineation, categorization or prediction. ML techniques have been traditionally applied to large, highly-dimensional databases. Gliomas are a heterogeneous group of primary brain tumors, traditionally graded using histopathological features. Recently the World Health Organization proposed a novel grading system for gliomas incorporating molecular characteristics. We aimed to study whether ML could achieve accurate prognostication of 2-year mortality in a small, highly-dimensional database of glioma patients.

**Methods:** We applied three machine learning techniques: artificial neural networks (ANN), decision trees (DT), support vector machine (SVM), and classical logistic regression (LR) to a dataset consisting of 76 glioma patients of all grades. We compared the effect of applying the algorithms to the raw database, versus a database where only statistically significant features were included into the algorithmic inputs (feature selection).

**Results:** Raw input consisted of 21 variables, and achieved performance of (accuracy/AUC): 70.7%/0.70 for ANN, 68%/0.72 for SVM, 66.7%/0.64 for LR and 65%/0.70 for DT. Feature selected input consisted of 14 variables and achieved performance of 73.4%/0.75 for ANN, 73.3%/0.74 for SVM, 69.3%/0.73 for LR and 65.2%/0.63 for DT.

**Conclusions:** We demonstrate that these techniques can also be applied to small, yet highly-dimensional datasets. Our ML techniques achieved reasonable performance compared to similar studies in the literature. Though local databases may be small versus larger cancer repositories, we demonstrate that ML techniques can still be applied to their analysis, though traditional statistical methods are of similar benefit.

## Introduction

Gliomas are a heterogeneous class of tumors comprising approximately 30% of all brain malignancies^1^. Previously, the World Health Organization (WHO) grading system stratified them by histological origin (i.e. astrocytoma, oligodendroglioma, mixed oligoastrocytoma and ependymoma), with additional grading (I-IV) according to pathological features of aggression. In 2016, the WHO presented a novel classification system, with incorporation of molecular biomarkers including IDH1/IDH2 mutations^2^, MGMT methylation^3^, p53 and PTEN deletion^4,5^, EGFR amplification^6^, 1p/19q deletions^7,8^, 9p(16q) deletions^9^ and Ki67 index^10^. The phenotypic expression of these markers by a glioma carry unique prognostic^11^ and therapeutic implications^6,7,11–14^. Moreover, the prognostic implications of the relationship between a tumor possessing >1 molecular marker with a patients’ baseline clinical and demographic status is not fully understood^15,16^. Existing prognostic systems separate patients into “low-grade” (i.e. WHO I,II) or “high-grade” (i.e. WHO III,IV) groups, and incorporate additional clinical features such as performance status, age and tumor size^13,17–21^ into their stratifications. Though some newer studies have incorporated limited molecular classification features^22^, it is clear that older prognostic indices are likely to become obsolete in the “molecular medicine” era.

Machine learning (ML) is a subset of computer science, whereby a computer algorithm “learns” from prior experience. Using specified training data with known input and output values, the ML algorithm is able to devise a set of rules which can be used as predictors for novel data with similar input characteristics^23^. Previously a human investigator would have to approach data collection and analysis using a set of *a priori* assumptions to prevent the burden of collecting data irrelevant to their hypothesis. The risk of this approach is that potentially meaningful trends caused by disregarded variables go unnoticed. ML lends itself naturally to trend-delineation in large, unprocessed datasets^24^. It can also be used for clinical prediction using known inputs and desired outputs (e.g. mortality). Moreover, when implemented in a local database, ML-derived prognosticators may take-into account unique features of the local population and treatment infrastructure, making them potentially more useful than evidence from non-contiguous populations. Local databases may be considerably smaller than large-scale cancer repositories, limiting their “academic study” in context of the literature, but potentially providing the local clinician with a wealth of meaningful clinical information.

Bearing these factors in mind, we aimed to apply a selection of ML algorithms to a database of 76 glioma cases in order to devise a 2-year mortality predictor. The complex histological and molecular pathological features of gliomas, combined with a series of clinical prognosticators, such as performance status, age and treatment techniques^25^, make them an ideal multi-dimensional application for machine learning techniques. Additionally, due to our database’ characteristics we aimed to compare the performance of ML algorithms using an unprocessed dataset with a dataset where only statistically-significant variables had been pre-selected.

## Machine Learning Methods

### Logistic Regression

Logistic regression (LR) **(Figure 1A)** is a traditional statistical method used for binary classification and has been adopted as a basic ML model. It differs from linear regression **(Figure 1B)** as it uses a sinusoidal curve, delineating a boundary between two categories. Like linear regression, the logarithmic function is derived from weighted transformation of the categorical data points. The regression function thus categorizes novel inputs into one of two categories based upon what side of the line its coordinates fall upon.

**Figure 1A-E.**
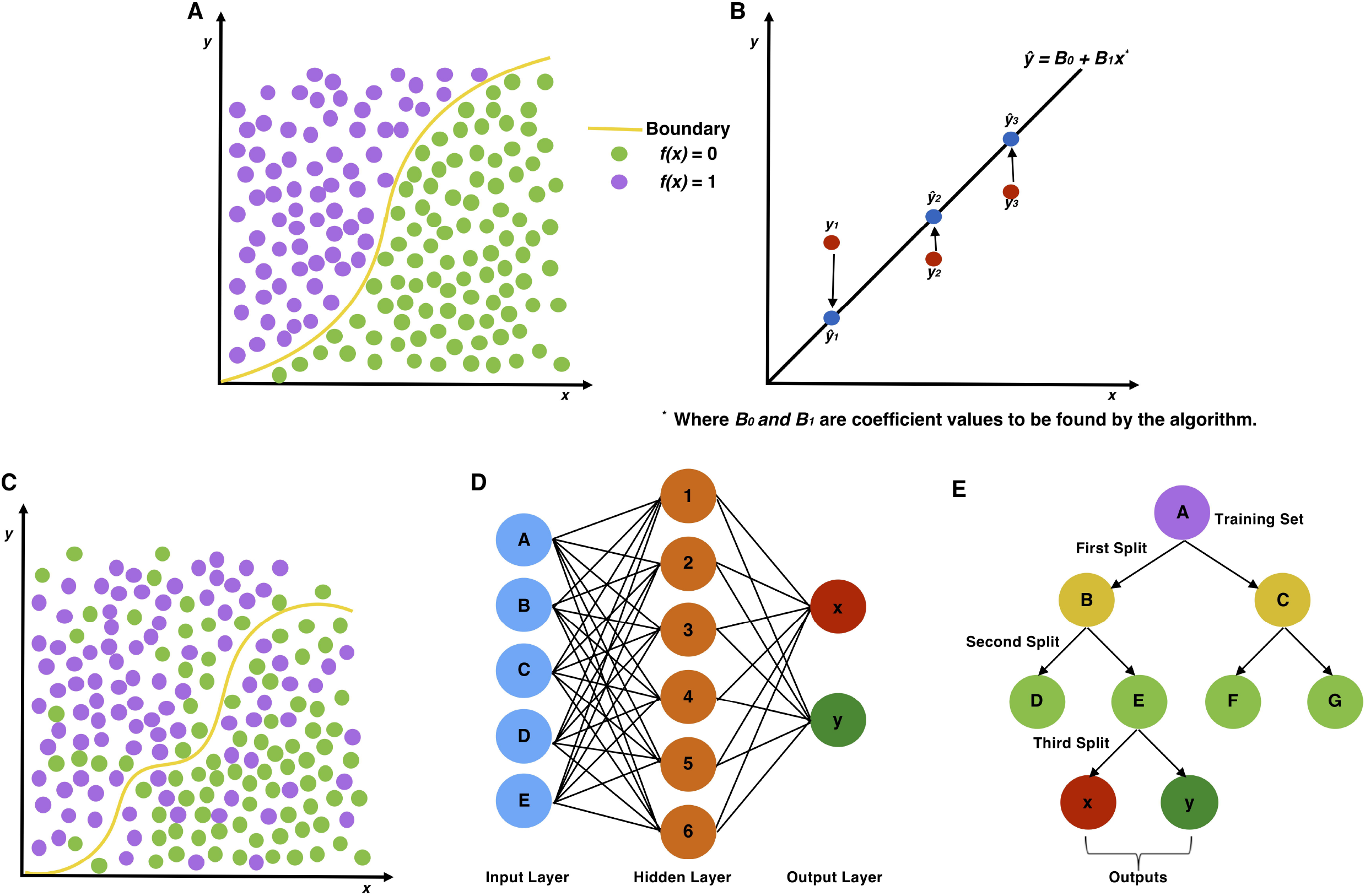
Graphical representation of traditional statistical approaches to regression, with logistic (A) and linear regression (B) on the top row. Bottom row demonstrates graphically ML approaches, with SVM (C), ANN (D), DT (E) approaches on the bottom row.

### Support Vector Machines

Support vector machines (SVM) **(Figure 1C)** are based upon the logistic regression method and assign training examples to one of two categories, with a bisecting “hyperplane” separating the data-points. Unlike logistic regression, however, the optimal hyperplane bisects the points representing the largest separation between the two categories, and its shape may not be defined by a simple function. The algorithm is tasked with finding the data points (“support vectors”) defining the hyperplane and derivative line coefficients. The function can then categorize novel input values into groups falling on either side of the hyperplane, similar to logistic regression.

### Artificial Neural Networks

Artificial neural networks (ANN) **(Figure 1D)** are so called because they are based upon the layer-like histological stratification of neurons. The input and output values represent the most superficial, but opposing layers of the network, while the inner “hidden” layers consist of successive transformations of the input values. ANNs can be used for clinical prediction algorithms by using a set of training data consisting of certain input values, with known output values. The algorithm therefore learns from the training set by progressive transformation of initial inputs. Values of these transformed inputs are then used by the model to predict output values.

### Decision Trees

Decision tree (DT) **(Figure 1E)** algorithms split data into binary categories using progressive iterations. ML algorithms aim to find optimal features at which to perform data splitting, creating a “branching-tree” shaped diagram. Each node represents a point at which the data is split, and the “leaves” at the end of the tree are the output variables. As the method involves binary classification, categorical data is preferred, while non-categorical data is preferably discretized prior to input.

## Methods

### Study Population

Our study population consisted of 76 patients (40 females, 36 males) with WHO grade I-IV gliomas, presenting to the neurosurgical oncology service at **[Redacted]** from 2009-2017. At the end of the 2-year followup period, 52 patients were alive, while 24 had perished. Mean age for the whole population at diagnosis was 47.3 years (SD 16.8). Interventions included total or subtotal resection (as stated by the operating surgeon), stereotactic biopsy, gamma knife therapy or no intervention. Other information collected included radiological maximum tumor diameter (cm), CNS tumor location (lobe), pre-operative and postoperative Eastern Cooperative Oncology Group (ECOG) performance status (0-5), subsequent chemotherapy, radiotherapy or vaccine therapy, or more than 1 surgical intervention. Surgical histopathology data included the presence of EGFR amplification, PTEN deletion, p53 mutation, 1p deletion, 19q deletion, 9p (p16) deletion, IDH 1/IDH 2 mutations, MGMT methylation, and Ki 67 proliferation index*.

### Ethical Considerations

This study was approved by the Institutional Review Board at the **[Redacted]**. All subjects consented for their de-identifiable data to be used in this study.

### Study Design

Due to the relatively small number of subjects in our database (n = 76), and the high dimensionality of the data, with 21 variables, we adopted two approaches to ML for this population **(Figure 2)**. The first was to apply the algorithms to the raw dataset, for which input variables had not been pre-selected. The second was to apply *χ*^2^ (for categorical variables) and independent samples *T*-tests (for continuous variables) to the dataset, as outlined by Oermann et al. (2013)^26^. As this involved a number of independent statistical tests, Bonferroni correction was applied subsequently. 14 variables were thus identified for which there was a significant difference between subjects who survived 2-years and those who did not. Non-significant variables were excluded from the input.

**Figure 2.**
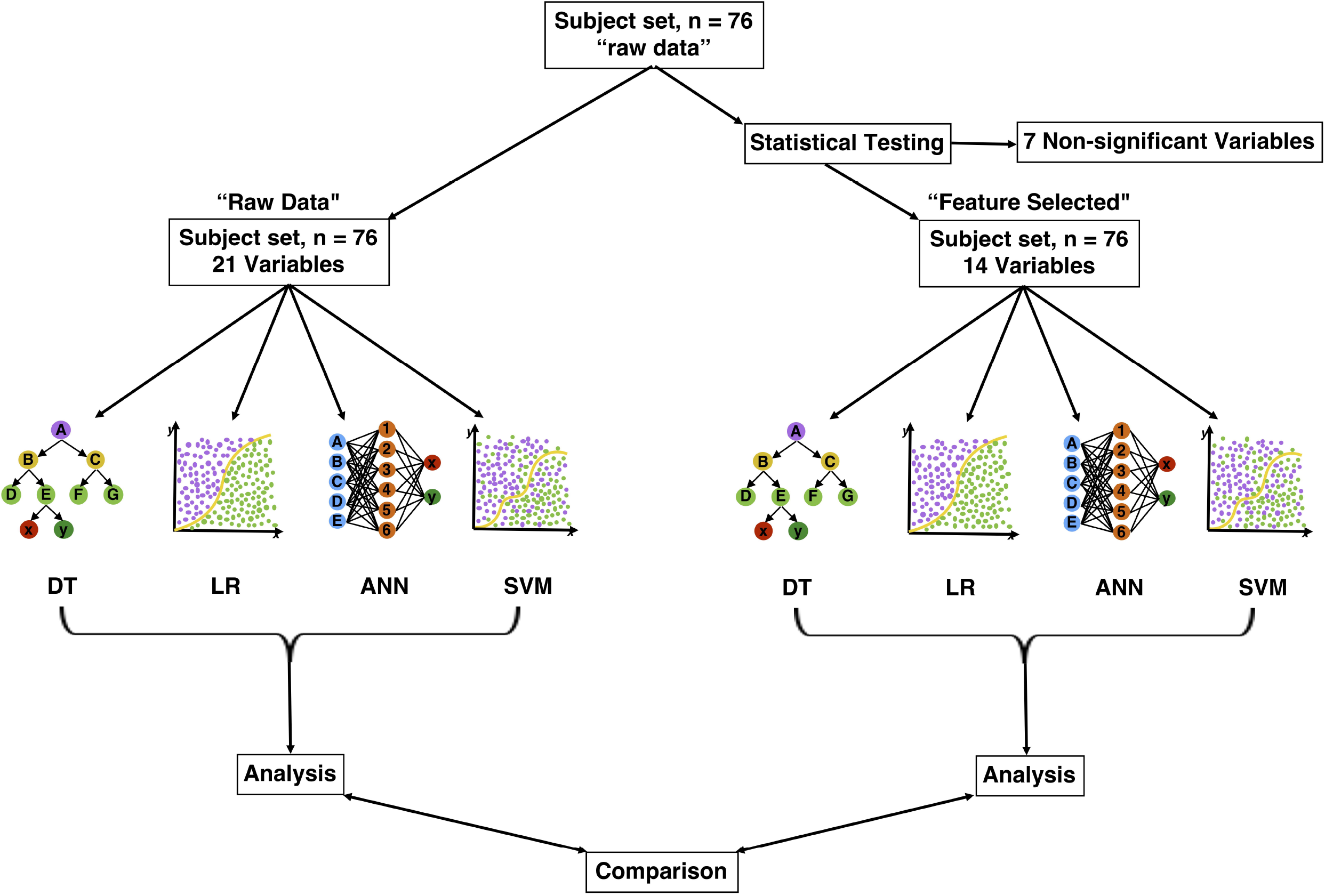
Diagram of study design demonstrating both approaches to ML. Left is the raw approach, right demonstrates initial statistical testing of variable significance prior to ML input. Outputs (performance) were analyzed independently and comparatively.

### Data Collection, Information Encoding and Dataset Splitting

The raw data was collected using Microsoft Excel (Microsoft Corporation, Seattle, U.S.A.), and the data was parsed using Python 2.7 programming language, using a custom written code. We used binary notation for ordinal variables (i.e. ‘yes’ = 1, ‘no’ = 0). Categorical and continuous variables were scaled (e.g. for Ki 67 index and age at diagnosis) to values between 0-1. The continuous variables were age, maximum tumor diameter and Ki 67 index. Categorical variables were total resection, performance status (ECOG), lobe/area of brain affected, WHO grade. There were 3 subjects whose surgical pathology results were unavailable. Instead of discarding these from analysis, we assigned a value of 0.5 for each variable (e.g. IDH1/2, PTEN etc.) All the features were then normalized using a normal vector.

### ML Algorithms

All ML and logistic regression models were imported from the SciKitLearn^27^ library. All models were run 15 times for each model. Each cycle consisted of a training and testing stage, where the dataset was repetitively partitioned. Per cycle, the same subject was not used for both training and testing. The number of dead and alive participants in the training and testing sets did vary between cycles, however.

### Artificial Neural Network Method

Our ANN method utilized a single layer of neurons between the input and output layers. The intermediate layer contained 100 neurons, each with a mini-batch size of 5. The network was trained using 1000 epochs, using an Adam optimizer^28^ with a default 0.001 learning rate. Briefly, the Adam optimizer is an algorithm for first-order gradient-based optimization, which is an extension to stochastic gradient descent.

### Decision Tree Method

The criteria used to split each node was determined by the Gini index^29^, a standard measure of information gain in DT applications^30^. This represents a more intuitive approach than randomly selecting criteria at which to split data. The minimum number of samples for each leaf was 1, while the minimum number of samples to split a node was 2.

### Support Vector Machine Method

Our SVM model used a radial basis function Kernel, with a C-penalty parameter of 100 and a Gamma value of 0.1.

### Logistic Regression

The penalization parameter used was l2 norm. The C parameter was 150.0 and the optimization algorithm used was coordinate descent.

### Data Processing

The averaged output values from the 15 cycles were then tabulated into standardized 4×4 confusion matrices and the sensitivity, specificity, positive likelihood ratio (PLR), negative likelihood ratio (NLR), positive predictive value (PPV), negative predictive value (NPV) and overall accuracy were calculated. All probabilities were calculated to 95% certainty. Receiver operating curves (ROC) and area under the ROC (AUROC) were additionally calculated and tabulated using the “roc_curve” model imported from the SciKitLearn toolbox. To optimize comparison between accuracy (percentages) and AUC (ratio), we multiplied AUC results by 100.

## Results

### Comparison of Diagnostic Performance

For raw data the ANN method performed best in terms of sensitivity (81.54%), followed by SVM (79.31%), LR (76.75%) and DT (73.65%) methods. Using a feature-selected dataset, sensitivity decreased for DT (68.93%), ANN (78.39%) and LR (74.26%), but increased slightly for SVM (80.54%). Using a feature-selected dataset, the specificity of all algorithms increased for all methods, with ANN performance showing the biggest increase (+11.62%) and DT showing the smallest (+7.56%). Using a feature-selected versus a raw dataset, all methods demonstrated a performance increase in terms of PPV (SVM = +7.69%; ANN = +7.08%; LR = +7.03%; DT = +5.87%), while all (DT = −6.21%; ANN = −3.79%; LR = −3.37%) but SVM (+0.54%) demonstrated a decrease in NPV performance. Likewise, ANN (+0.42), SVM (+0.36), LR (+0.28), DT (+0.14) demonstrated an increase in PLR performance using a feature-selected dataset. In terms of NLR, all methods (SVM = −0.10; LR = −0.04; ANN = −0.02) aside from DT (+0.55) demonstrated a decrease in NLR prediction. All methods demonstrated an increase in accuracy using the feature-selected dataset (SVM = +5.38%; ANN = +2.71%; LR = +2.62%; DT = +0.17%). Finally, censoring increased performance, as represented by AUC for all methods (LR = +8.58%; ANN = +6.21%; SVM = +2.83%) aside from DT, which demonstrated a decrease in AUC (−7.54%)**(Figure 3)^†^**.

**Figure 3.**
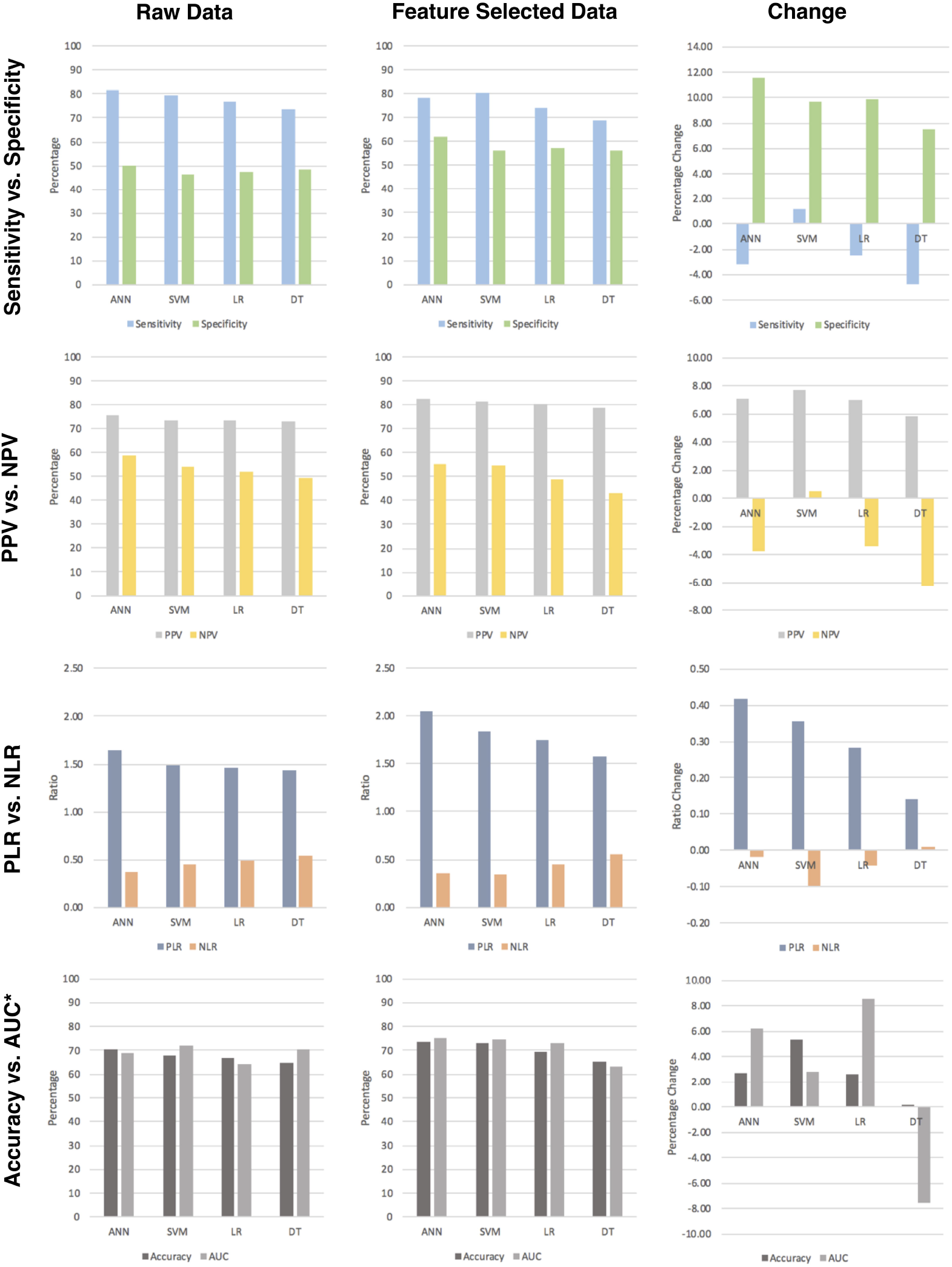
A 4×3 array of figures demonstrating the algorithm performance using both feature-selected (far left column) and feature-selected (middle column) approaches. Change in performance between raw and feature-selected data is demonstrated in the far-right column. First row shows sensitivity vs. specificity performance; second row shows PPV vs. NPV performance; third row shows PLR vs. NLR performance; fourth row shows accuracy vs. AUC performance. *AUC metrics have been scaled to 100 to correlate with accuracy.

### Receiver Operating Curves and Confidence Intervals

When comparing the ROC curve performance to that of *y*=*x*, with an area of 0.5 (50), the SVM (AUC = 71.88) demonstrated the best performance, followed by DT (AUC = 70.54), ANN (AUC = 69.19) and LR (AUC = 64.29). Though these were higher than 0.5, the 95% confidence intervals (C.I.) for both the ANN (C.I. = 49.86 - 88.52) and LR (C.I. = 43.63 − 84.95) both included 50, indicating non-significance. Even though the SVM (C.I. = 53.40 − 90.36) and DT (C.I. = 51.62 − 89.46) algorithms had lower C.I. values >50, these were only marginally higher than this boundary. The feature-selected datasets, provided a performance increase for all but the DT algorithms, which demonstrated a decrease in AUC value. The performance benefit was indicated by higher AUC values, with ANN (AUC = 75.40) performing best, followed by SVM (AUC = 74.71), LR (AUC = 72.87) and DT (AUC = 63.00). Using feature-selected data also yielded overall narrower 95% C.I., with all methods aside from DT demonstrating at least a 10 unit increase of lower C.I. boundary above 50, indicating significance over random guessing and use of raw data. Nevertheless, for both feature-selected and raw data, none of the ML methods demonstrated significant performance improvement versus LR, nor over one another **(Figure 4)^‡^**.

**Figure 4.**
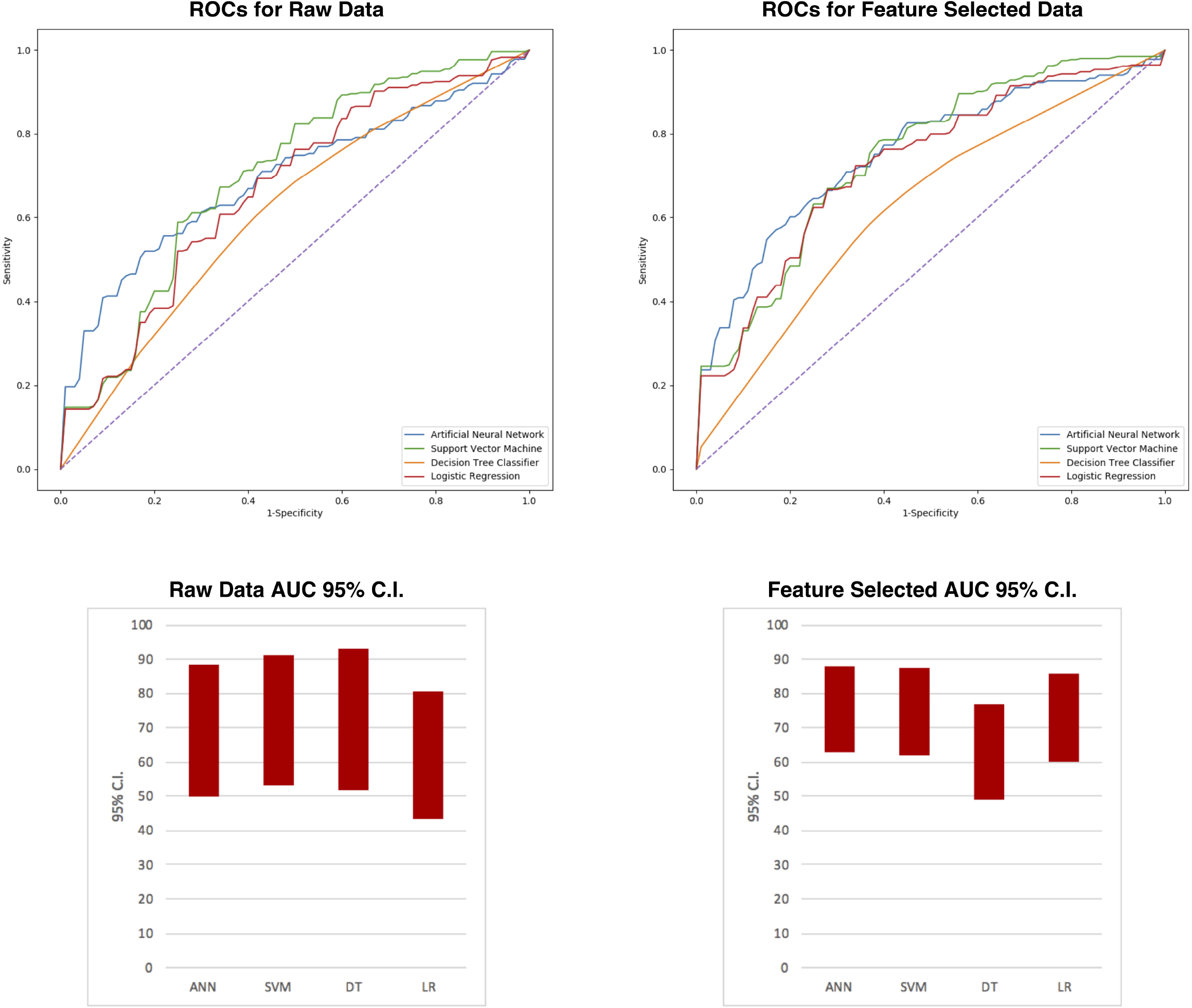
Reciever operating curves (ROC) for raw (left) and feature-selected (data) right. Lines are color coordinated using figure legend in bottom right corner of graph. Perforated diagonal line is y=x, with an AUC of 0.5 (random guessing). The performance increase is discerned by the increased distance between all curves and that of y=x. Lower right column chart demonstrates the scaled 95% confidence intervals of the AUCs calculated for each ML method. ANN and LR methods were not statistically difference from 0.5. SVM and DT methods were better than random guessing, however their statistical significance was weak. Performance of algorithms using raw datasets can be compared with the chart in the lower right corner. These are the 95% confidence intervals for all ML methods, and can be concluded to be not only further away from 0.5 but also narrower, indicating greater significance and less variation.

## Discussion

We have successfully demonstrated the application of 3 machine learning techniques and a ML-implemented logistic regression technique to a database of 76 glioma patients of all stages, molecular phenotypes and with heterogeneous clinical characteristics. Relative to older prognostic indices, which do not incorporate molecular features, our study involves considerably fewer subjects. We accomplished our goal of applying ML techniques to this database with a relatively low subject number/variable ratio, and furthermore we demonstrate that machine learning can be applied with a reasonable level of confidence to make prognostic inferences from this data.

### Comparison to Similar Studies

In the neuro-oncology literature, much focus of ML application has been directed towards discernment of characteristic imaging magnetic resonance imaging (MRI) characteristics of CNS tumors (discussed later). Only one non-imaging focussed study has utilized ML for glioma outcome prediction^31^, while the study of Oermann et al. (2013) utilized a similar methodology for cerebral metastasis prognostication. The study by Malhotra et al. (2016) applied a novel data mining algorithm to extract relevant features pertaining to treatment and molecular patterns in a database of 300 newly diagnosed glioblastoma multiforme cases. The ML component of their study involved the extraction of relevant treatment and pathological features, which were then classified and subjected to classical statistical methods for prognostication. This is in effect the opposite approach to our method using feature-selected data, as the authors utilized data mining to extract relevant features, while we conducted statistical tests of significance for feature-selection prior to ML input. They achieved maximal C-values of 0.85 using LR and 0.84 using Cox multivariate regression. The study of Oermann et al. (2013), though pertaining to cerebral metastases versus gliomas, utilized a similar methodology to our feature-selected approach to prognosticate 1-year survival in a total of 196 patients. In this study, the pooled voting results of 5 independent ANNs (AUC = 84%) significantly outperformed traditional LR methods (AUC = 75%). Further, they found that ML techniques were more accurate at predicting 1-year survival than 2 traditional prognostic indices. As our study used data from gliomas of all stages, we did not compare our results to existing prognostic indices which specifically separate patients into low and high-grade categories. Though even using a feature-selected approach, our best performing algorithm (which was coincidentally also an ANN) achieved approximately 10-unit lower AUC metric than their ANN approach. We suspect that this is due to two reasons: Firstly, their training set consisted of 98 patients, which was over twice the size of our training set of 40, offering more examples to learn from. Secondly, their method only utilized 6 input variables, versus 21 for our raw approach and 14 for our feature-selected approach. It is therefore likely that increased proportionality of subjects-to-variables in their dataset also enhanced predictive performance of the ML algorithm. From this, it is apparent that smaller datasets may require pre-censoring prior to ML application if predictive performance is to be maximized. Secondly, we cannot conclude that for small, highly-dimensional datasets ML approaches including ANN, DT or SVM offer any significant performance advantage over traditional LR methods. Nevertheless, we achieved reasonably good predictive metrics using pre-input variable censoring using all ML approaches.

### Future Directions

ML algorithms have been intuitively applied to “data-rich” MRI sequences in an effort to quantitatively discern characteristic imaging features of gliomas, due to differences between tumor area and normal brain^32–36^. These methods have yielded ability to discern imaging features indicating the presence of MGMT methylation^35^, IDH1 mutation^37–39^ and 1p/19q co-deletion^38^. These breakthroughs potentially allow for the non-invasive identification^40,41^ and even prognostication^41,42^ of gliomas according to imaging characteristics alone. It is not unreasonable to suggest that the next generation of prognostic indices will be derived from a combination of clinical database mining techniques, such as our present study, with novel techniques of image-based ML. This will represent a substantial step forward, as previous prognostic systems relied upon invasive methods for definitive diagnosis, prognostication and treatment plans. It may also permit clinicians to prognose non-invasively and with high-accuracy, the clinical course of low-grade tumors.

Another potential future direction for ML in neuro-oncology is the creation of a local repository of cases that can be used much in the same way as the study populations utilized for developing previous prognostication systems. Depending upon the integrity and scale of the localized database, predictions can be made with reasonable accuracy, as we have demonstrated in our present study. The information contained in a localized oncology repository may be of greater benefit to the clinician than data from a non-contiguous population, as localized population characteristics and demographics can be discerned and incorporated into individually-tailored therapeutic approaches.

### Limitations

Though we achieved acceptable predictive performance using feature-selected data, our study has highlighted potential difficulties of ML application to smaller, highly-dimensional clinical databases. Pre-censoring of data may optimize ML algorithms in studies using a smaller subject-set, however, the censoring of particular variables may result in weaker trends going unnoticed. We also anticipate that the predictive accuracy and AUC would improve by increasing the number of subjects included in the training set. Despite this, our study used a training set less than half the size of that of Oermann et al. (2013) yet achieved only slightly weaker prognostic performance. An alternative to the manual testing of variables for significance is principal component analysis^43^, which is an entirely ML-based method of reducing data-set dimensionality, thus reducing scope for human error or bias.

### Conclusions

Though the “data mining” of “big data” is popular phrase amongst both the medical and lay media, we have demonstrated that ML techniques are applicable to small yet highly-dimensional datasets. As the clinical approaches to gliomas are beginning to adapt to the molecular-medicine era, the small size of a local database does not provide a barrier to the implementation of ML techniques for prognostication purposes. Though our study was purely academic, it demonstrates the potential for ML to provide meaningful insight into the diagnosis and treatment of these heterogeneous tumors at a local level.

## Acknowledgements

We would like to thank Yue-Fang Chang PhD, at the University of Pittsburgh for her assistance with this project.

## References

1. Goodenberger ML, Jenkins RB. Genetics of adult glioma. Cancer Genet. 2012;205(12):613–621. doi:10.1016/j.cancergen.2012.10.009

2. Yan H, Parsons DW, Jin G, et al. IDH1 and IDH2 Mutations in Gliomas. N Engl J Med. 2009;360(8):765–773. doi:10.1056/NEJMoa0808710

3. Costello JF, Futscher BW, Tano K, Graunke DM, Pieper RO. Graded methylation in the promoter and body of the O6-methylguanine DNA methyltransferase (MGMT) gene correlates with MGMT expression in human glioma cells. J Biol Chem. 1994;269(25):17228–17237.

4. Zheng H, Ying H, Yan H, et al. p53 and Pten control neural and glioma stem/progenitor cell renewal and differentiation. Nature. 2008;455(7216):1129–1133. doi:10.1038/nature07443

5. Broniscer A, Baker SJ, West AN, et al. Clinical and molecular characteristics of malignant transformation of low-grade glioma in children. J Clin Oncol. 2007;25(6):682–689.

6. Smith JS, Tachibana I, Passe SM, et al. PTEN Mutation, EGFR Amplification, and Outcome in Patients With Anaplastic Astrocytoma and Glioblastoma Multiforme. JNCI J Natl Cancer Inst. 2001;93(16):1246–1256. doi:10.1093/jnci/93.16.1246

7. Eckel-Passow JE, Lachance DH, Molinaro AM, et al. Glioma Groups Based on 1p/19q, IDH, and TERT Promoter Mutations in Tumors. N Engl J Med. 2015;372(26):2499–2508. doi:10.1056/NEJMoa1407279

8. Kaloshi G, Benouaich-Amiel A, Diakite F, et al. Temozolomide for low-grade gliomas. Neurology. 2007;68(21):1831–1836.

9. Houillier C, Mokhtari K, Carpentier C, et al. Chromosome 9p and 10q losses predict unfavorable outcome in low-grade gliomas. Neuro-Oncol. 2010;12(1):2–6. doi:10.1093/neuonc/nop002

10. Preusser M, Hoeftberger R, Woehrer A, et al. Prognostic value of Ki67 index in anaplastic oligodendroglial tumours – a translational study of the European Organization for Research and Treatment of Cancer Brain Tumor Group. Histopathology. 60(6):885–894. doi:10.1111/j.1365-2559.2011.04134.x

11. Phillips HS, Kharbanda S, Chen R, et al. Molecular subclasses of high-grade glioma predict prognosis, delineate a pattern of disease progression, and resemble stages in neurogenesis. Cancer Cell. 2006;9(3):157–173. doi:10.1016/j.ccr.2006.02.019

12. Metellus P, Coulibaly B, Colin C, et al. Absence of *IDH* mutation identifies a novel radiologic and molecular subtype of WHO grade II gliomas with dismal prognosis. Acta Neuropathol (Berl). 2010;120(6):719–729. doi:10.1007/s00401-010-0777-8

13. Stupp R, Hegi ME, Mason WP, et al. Effects of radiotherapy with concomitant and adjuvant temozolomide versus radiotherapy alone on survival in glioblastoma in a randomised phase III study: 5-year analysis of the EORTC-NCIC trial. Lancet Oncol. 2009;10(5):459–466. doi:10.1016/S1470-2045(09)70025-7

14. Leu S, von Felten S, Frank S, et al. IDH/MGMT-driven molecular classification of low-grade glioma is a strong predictor for long-term survival. Neuro-Oncol. 2013;15(4):469–479. doi:10.1093/neuonc/nos317

15. Figarella-Branger D, Bouvier C, Paula AM de, et al. Molecular genetics of adult grade II gliomas: towards a comprehensive tumor classification system. J Neurooncol. 2012;110(2):205–213. doi:10.1007/s11060-012-0953-x

16. Zalatimo O, Zoccoli CM, Patel A, Weston CL, Glantz M. Impact of Genetic Targets on Primary Brain Tumor Therapy: What’s Ready for Prime Time? In: Impact of Genetic Targets on Cancer Therapy. Advances in Experimental Medicine and Biology. Springer, New York, NY; 2013:267–289. doi:10.1007/978-1-4614-6176-0_12

17. Council MR. Prognostic factors for high-grade malignant glioma: Development of a prognostic index. J Neurooncol. 1990;9(1):47–55. doi:10.1007/BF00167068

18. Gorlia T, Wu W, Wang M, et al. New validated prognostic models and prognostic calculators in patients with low-grade gliomas diagnosed by central pathology review: a pooled analysis of EORTC/RTOG/NCCTG phase III clinical trials. Neuro-Oncol. 2013;15(11):1568–1579. doi:10.1093/neuonc/not117

19. van den Bent MJ, Afra D, de Witte O, et al. Long-term efficacy of early versus delayed radiotherapy for low-grade astrocytoma and oligodendroglioma in adults: the EORTC 22845 randomised trial. Lancet Lond Engl. 2005;366(9490):985–990. doi:10.1016/S0140-6736(05)67070-5

20. Mirimanoff R-O, Gorlia T, Mason W, et al. Radiotherapy and temozolomide for newly diagnosed glioblastoma: recursive partitioning analysis of the EORTC 26981/22981-NCIC CE3 phase III randomized trial. J Clin Oncol. 2006;24(16):2563–2569.

21. Karim ABMF, Maat B, Hatlevoll R, et al. A randomized trial on dose-response in radiation therapy of low-grade cerebral glioma: European Organization for Research and Treatment of Cancer (EORTC) study 22844. Int J Radiat Oncol • Biol • Phys. 1996;36(3):549–556. doi:10.1016/S0360-3016(96)00352-5

22. Daniels TB, Brown PD, Felten SJ, et al. Validation of EORTC Prognostic Factors for Adults With Low-Grade Glioma: A Report Using Intergroup 86-72-51. Int J Radiat Oncol • Biol • Phys. 2011;81(1):218–224. doi:10.1016/j.ijrobp.2010.05.003

23. Obermeyer Z, Emanuel EJ. Predicting the Future — Big Data, Machine Learning, and Clinical Medicine. N Engl J Med. 2016;375(13):1216–1219. doi:10.1056/NEJMp1606181

24. Cruz JA, Wishart DS. Applications of Machine Learning in Cancer Prediction and Prognosis. Cancer Inform. 2006;2:117693510600200030. doi:10.1177/117693510600200030

25. Jeremic B, Milicic B, Grujicic D, Dagovic A, Aleksandrovic J, Nikolic N. Clinical Prognostic Factors in Patients With Malignant Glioma Treated With Combined Modality Approach. Am J Clin Oncol. 2004;27(2):195. doi:10.1097/01.coc.0000055059.97106.15

26. Oermann EK, Kress M-AS, Collins BT, et al. Predicting Survival in Patients With Brain Metastases Treated With Radiosurgery Using Artificial Neural Networks. Neurosurgery. 2013;72(6):944–952. doi:10.1227/NEU.0b013e31828ea04b

27. Pedregosa F, Varoquaux G, Gramfort A, et al. Scikit-learn: Machine Learning in Python. J Mach Learn Res. 2011;12(Oct):2825–2830.

28. Kingma DP, Ba J. Adam: A Method for Stochastic Optimization. ArXiv14126980 Cs. December 2014. http://arxiv.org/abs/1412.6980. Accessed June 9, 2018.

29. Gastwirth JL. The Estimation of the Lorenz Curve and Gini Index. Rev Econ Stat. 1972;54(3):306–316. doi:10.2307/1937992

30. Raileanu LE, Stoffel K. Theoretical Comparison between the Gini Index and Information Gain Criteria. Ann Math Artif Intell. 2004;41(1):77–93. doi:10.1023/B:AMAI.0000018580.96245.c6

31. Malhotra K, Navathe SB, Chau DH, Hadjipanayis C, Sun J. Constraint based temporal event sequence mining for Glioblastoma survival prediction. J Biomed Inform. 2016;61:267–275. doi:10.1016/j.jbi.2016.03.020

32. Macyszyn L, Akbari H, Pisapia JM, et al. Imaging patterns predict patient survival and molecular subtype in glioblastoma via machine learning techniques. Neuro-Oncol. 2016;18(3):417–425. doi:10.1093/neuonc/nov127

33. Kickingereder P, Bonekamp D, Nowosielski M, et al. Radiogenomics of Glioblastoma: Machine Learning-based Classification of Molecular Characteristics by Using Multiparametric and Multiregional MR Imaging Features. Radiology. 2016;281(3):907–918. doi:10.1148/radiol.2016161382

34. Zacharaki EI, Wang S, Chawla S, et al. Classification of brain tumor type and grade using MRI texture and shape in a machine learning scheme. Magn Reson Med. 62(6):1609–1618. doi:10.1002/mrm.22147

35. Ahn SS, Shin N-Y, Chang JH, et al. Prediction of methylguanine methyltransferase promoter methylation in glioblastoma using dynamic contrast-enhanced magnetic resonance and diffusion tensor imaging. J Neurosurg. 2014;121(2):367–373. doi:10.3171/2014.5.JNS132279

36. Ranjith G, Parvathy R, Vikas V, Chandrasekharan K, Nair S. Machine learning methods for the classification of gliomas: Initial results using features extracted from MR spectroscopy. Neuroradiol J. 2015;28(2):106–111. doi:10.1177/1971400915576637

37. Yu J, Shi Z, Lian Y, et al. Noninvasive IDH1 mutation estimation based on a quantitative radiomics approach for grade II glioma. Eur Radiol. 2017;27(8):3509–3522. doi:10.1007/s00330-016-4653-3

38. Zhou H, Vallières M, Bai HX, et al. MRI features predict survival and molecular markers in diffuse lower-grade gliomas. Neuro-Oncol. 2017;19(6):862–870. doi:10.1093/neuonc/now256

39. Zhang B, Chang K, Ramkissoon S, et al. Multimodal MRI features predict isocitrate dehydrogenase genotype in high-grade gliomas. Neuro-Oncol. 2017;19(1):109–117. doi:10.1093/neuonc/now121

40. Wiestler B, Kluge A, Lukas M, et al. Multiparametric MRI-based differentiation of WHO grade II/III glioma and WHO grade IV glioblastoma. Sci Rep. 2016;6:35142. doi:10.1038/srep35142

41. Chang K, Zhang B, Guo X, et al. Multimodal imaging patterns predict survival in recurrent glioblastoma patients treated with bevacizumab. Neuro-Oncol. 2016;18(12):1680–1687. doi:10.1093/neuonc/now086

42. Emblem KE, Due-Tonnessen P, Hald JK, et al. Machine learning in preoperative glioma MRI: Survival associations by perfusion-based support vector machine outperforms traditional MRI. J Magn Reson Imaging. 40(1):47–54. doi:10.1002/jmri.24390

43. Ringnér M. What is principal component analysis? Nat Biotechnol. 2008;26(3):303–304. doi:10.1038/nbt0308-303

